## Supplemental Figures and Table for "Functional and structural deficiencies of Gemin5 variants associated with neurological disease"

**SUPPLEMENTARY FIGURES**

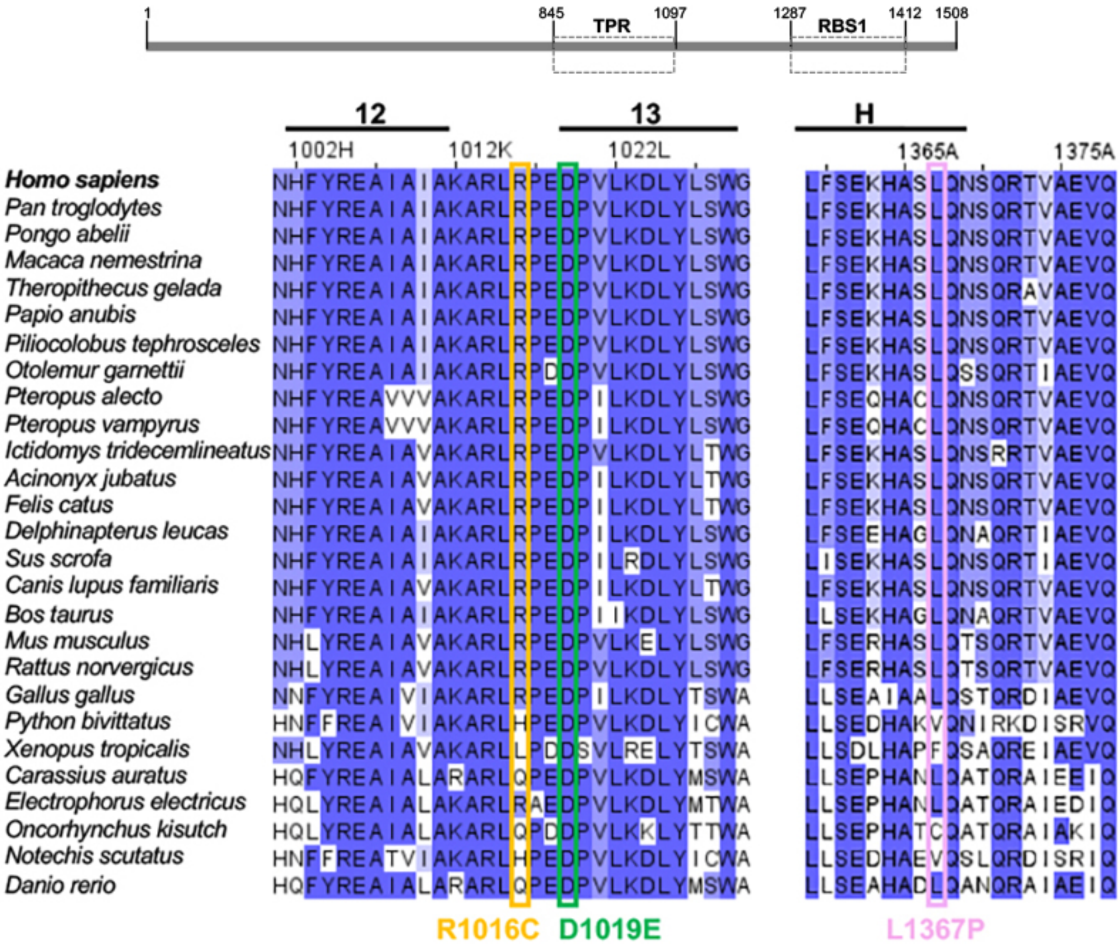


**Figure S1**. Alignment of Gemin5 sequences from 27 species spanning helices 12 and 13 of the TPR-like domain and a C-terminal helix of RBS1, colored according to their degree of conservation, from blue to white. Boxes orange, green and pink denote the sequence variability of R1016, D1019, and L1367, respectively.

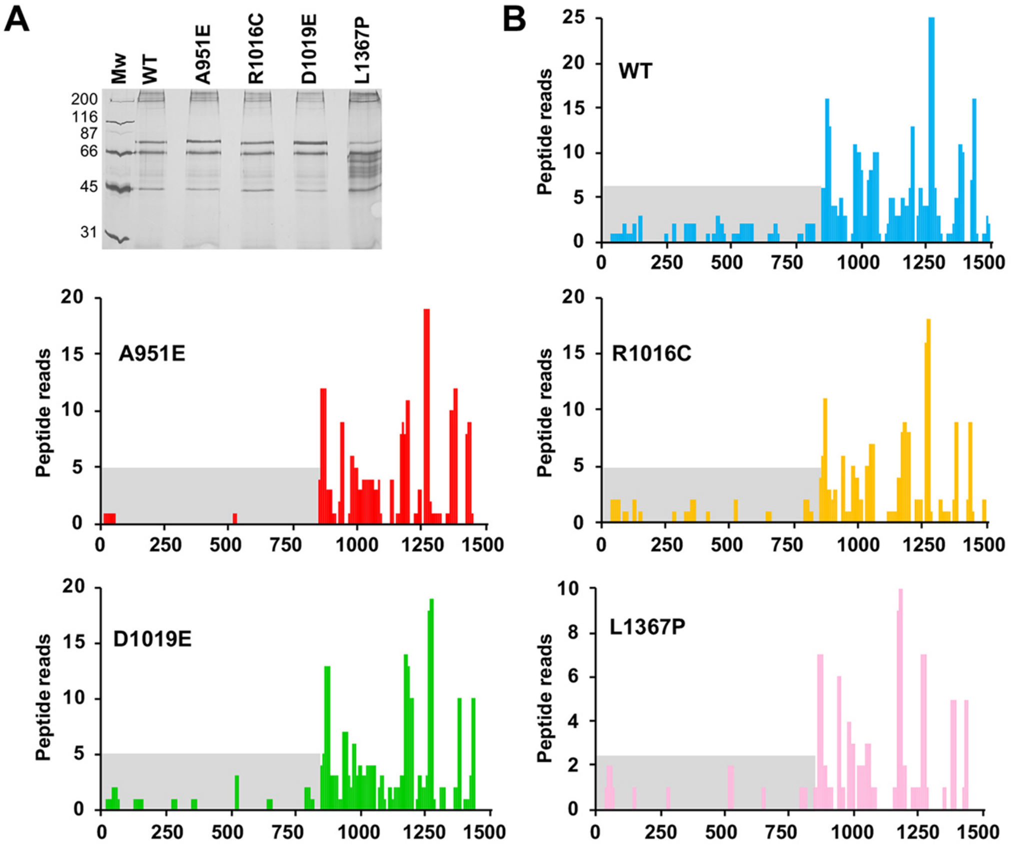


C

| TOTAL READS | WT 1 | WT 2 | A951E 1 | A951E 2 | R1016C 1 | R1016C 2 | D1019E 1 | D1019E 2 | L1367P 1 | L1367P 2 |
|---|---|---|---|---|---|---|---|---|---|---|
| 1-844 | 311 | 540 | 32 | 47 | 252 | 153 | 174 | 189 | 129 | 76 |
| 845-1508 | 1895 | 2624 | 1348 | 1833 | 1623 | 1184 | 1829 | 1918 | 857 | 418 |

**Figure S2. Gemin5 peptides identification in $G5_{845\text{-}1508}$-TAP samples. (A)** Silver stained gel loaded with TAP samples obtained with the indicated variants of $G5_{845\text{-}1508}$. **(B)** Graphs represent the number of peptide reads according to the amino acid position determined in one of the replicates of $G5_{845\text{-}1508}$-WT (blue), A951E (red), R1016C (yellow), D1019E (green), and L1367P (pink). The peptide reads corresponding to the N-terminal region of Gemin5 (residues 1-844) are highlighted in grey background. **(C)** Summary of the total reads obtained by LC-MS/MS for the region 1-844, and 845-1508 with two independent replicas.

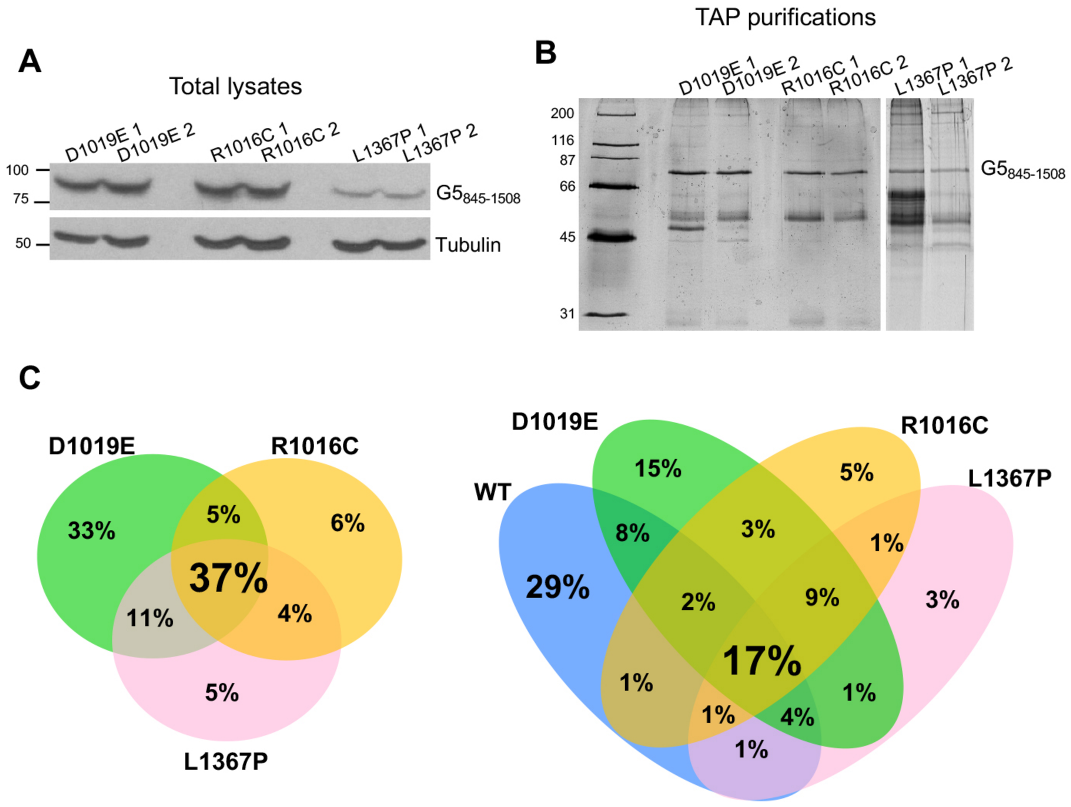


**Figure S3. Representative examples of protein purification for analysis of proteins associated with G5$_{845-1508}$. (A)** Immunodetection of G5$_{845-1508}$-TAP variants in total lysates of HEK293 transfected cells. The proteins were detected with anti-CBP and tubulin was used as loading control. **(B)** Silver stained gel loaded with two biological replicates for each G5$_{845-1508}$ variant subjected to proteomic analysis. **(C)** Venn diagram of proteins identified with the three Gemin5 clinical variants (left). Comparison of the protein-interactome of the three variants with the with the WT protein (right)

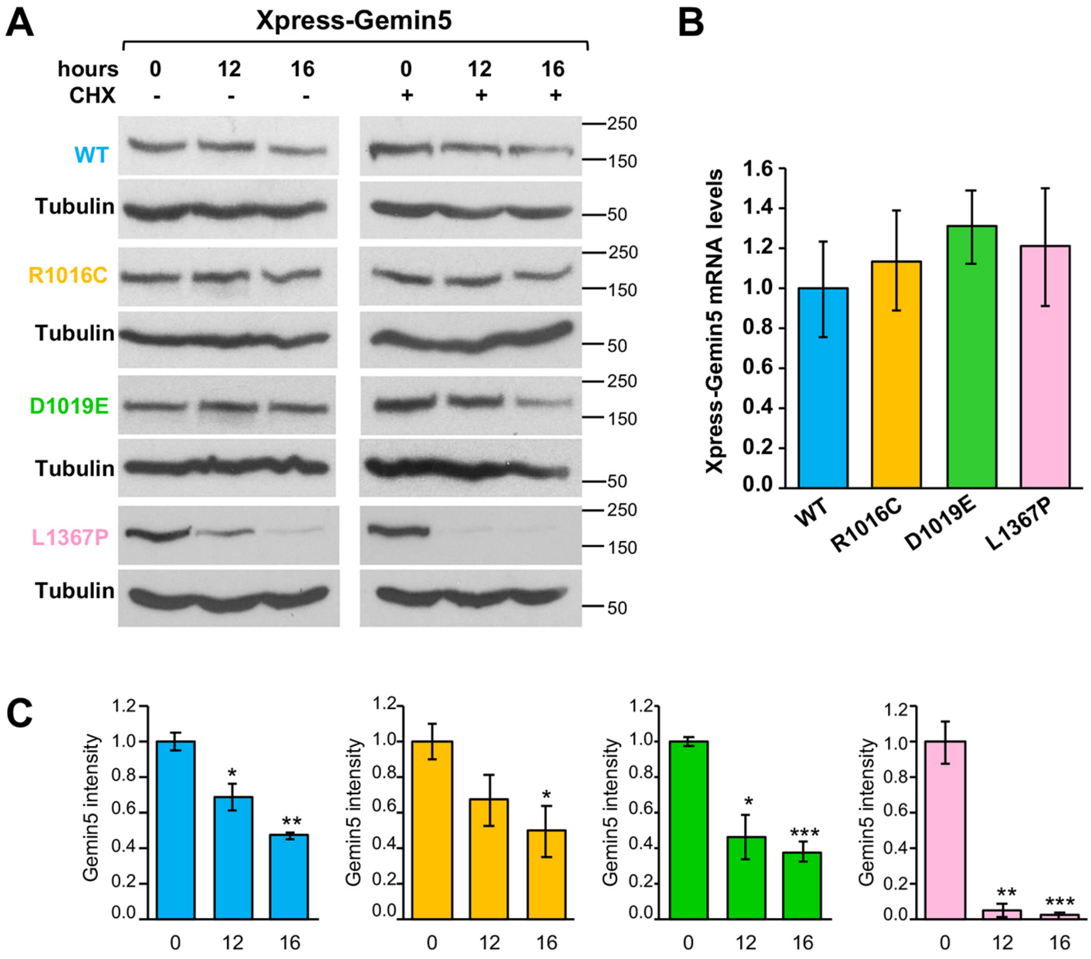


**Figure S4. Protein stability of Gemin5 carrying mutations R1016C, D1019E or L1367P. (A)** HEK293 cells expressing the wild type version of Xpress-Gemin5, side by side to the variants R1016C, D1019E or L1367P during 24 h were treated (+) or not (-) with cycloheximide (CHX) for additional 16 h. Samples were taken at 0, 12 and 16 h post CHX treatment. The intensity of each protein at the indicated time was determined by WB. **(B)** Steady-state analysis of Xpress-Gemin5 mRNA levels present in transfected cells at the time of harvesting determined by RTqPCR. **(C)** Values represent the protein intensity (mean ± SEM) of three independent experiments relatively to time 0. Asterisks denote *P*-values (**P* < 0.05, ***P* < 0.01, ****P* < 0.001).

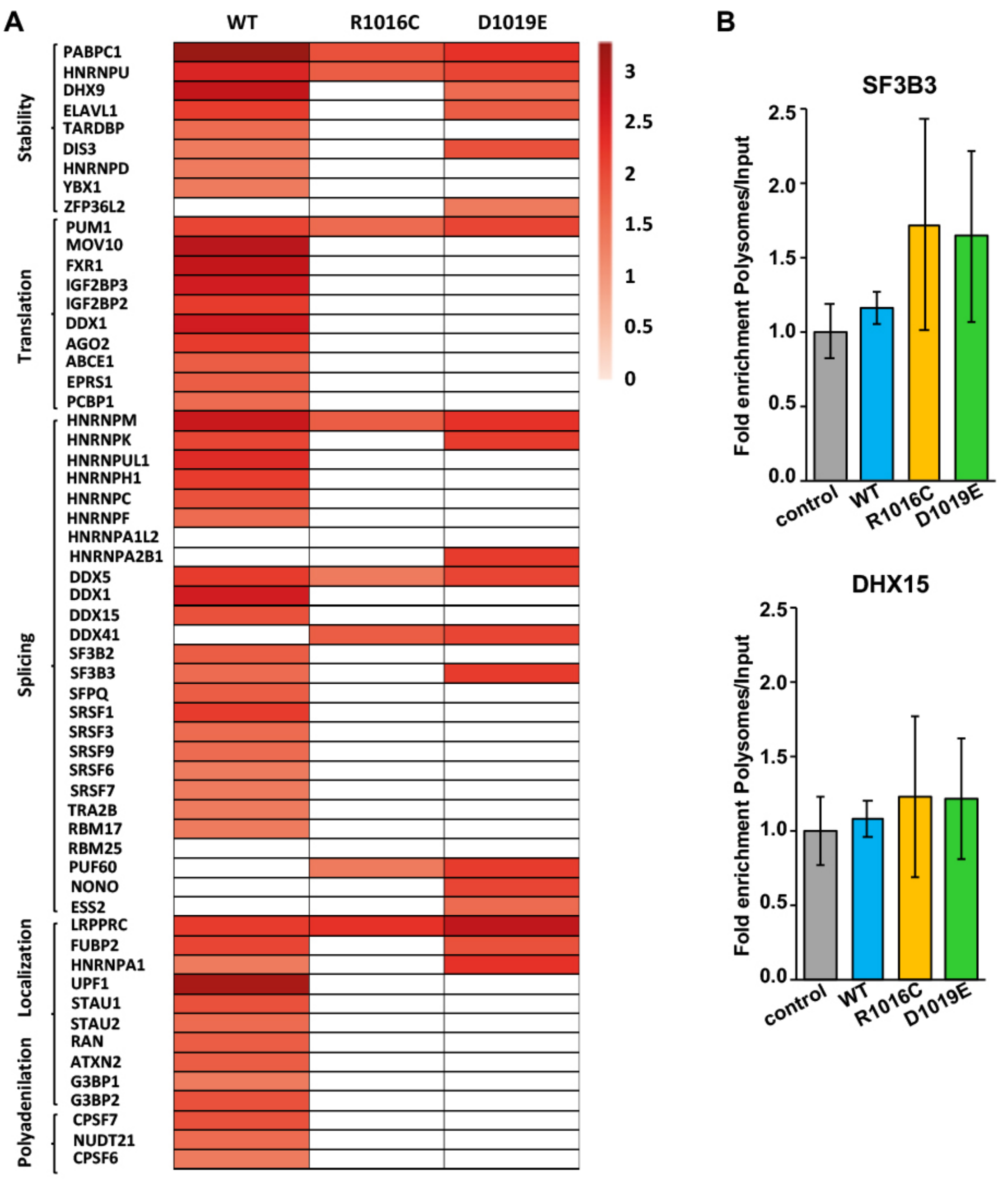


**Figure S5. (A)** Heat map representation of RNA-binding proteins associated to Gemin5 variants. Proteins involved in RNA stability, translation regulation, splicing, localization and polyadenylation identified by LC-MS/MS are depicted according to ln of peptides identified from dark red to white. **(B)** Histograms representing the fold enrichment in polysomes relative to the input samples of selected mRNAs (SF3B3 and DHX15).

**Table S1. Primers used in constructs and RTqPCR analysis.**

| Oligo | Sequence (5´- 3´) |
|---|---|
| G5-D1019E s | tcaggactggctcctccgggcgc |
| G5-D1019E as | gcgcccggaggagccagtcctga |
| G5-R1016C s | gtcctccgggcacagccgggcct |
| G5-R1016C as | aggcccggctgtgcccggaggac |
| G5-L1367P s | gttctctgtgagttttggggactggcatgcttttctg |
| G5-L1367P as | cagaaaagcatgccagtccccaaaactcacagagaac |
| 5'EcoRI-GST-RPS9 | aagaattcacatgccagtgg |
| 3'SalI-GST-RPS9 | atgtcgacttaatcctcctcctcgt |
| 5'BamHI-RPS26 | taggatccatgacaaagaaaagaagg |
| 3'SalI-GST-RPS26 | atgtcgacttacatgggctttgg |
| His-Xpress s | ggggttctcatcatcatcatc |
| His-Xpress as | gatccttatcgtcatcgtcgt |
| AGO2-s | ataaagctattgcgacccctg |
| AGO2-as | tcagatggacttccgtgc |
| PCBP1-s | attcgccggaattgactcca |
| PCBP1-as | atggtgagttcatgggtggt |
| SF3B3-s | agaaatgttccagcccaccc |
| SF3B3-as | acttctccaggttcaaggcg |
| DHX15-s | ggtgtgtggagtacatgcga |
| DHX15-as | cagcaactctctgagccaca |
